# Calcium induced N-terminal gating and pore collapse in connexin-46/50 gap junctions

**DOI:** 10.1101/2025.02.12.637955

**Authors:** Jonathan A. Flores, Susan E. O’Neill, Joshua M. Jarodsky, Steve L. Reichow

**Affiliations:** Department of Chemical Physiology and Biochemistry, Oregon Health and Science University, Portland, OR 97239, USA; Vollum Institute, Oregon Health and Science University, Portland, OR 97239, USA; Currently: Department of Molecular Biology and Genetics, Aarhus University, Aarhus, Denmark

**Keywords:** connexin, gap junction, cryo-EM, calcium regulation, channel gating, large-pore channel

## Abstract

Gap junctions facilitate electrical and metabolic coupling essential for tissue function. Under ischemic conditions (*e.g.,* heart attack or stroke), elevated intracellular calcium (Ca^2+^) levels uncouple these cell-to-cell communication pathways to protect healthy cells from cytotoxic signals. Using single-particle cryo-EM, we elucidate details of the Ca^2+^-induced gating mechanism of native connexin-46/50 (Cx46/50) gap junctions. The resolved structures reveal Ca^2+^ binding sites within the channel pore that alter the chemical environment of the permeation pathway and induce diverse occluded and gated states through N-terminal domain remodeling. Moreover, subunit rearrangements lead to pore collapse, enabling steric blockade by the N-terminal domains, reminiscent of the “iris model” of gating proposed over four decades ago. These findings unify and expand key elements of previous gating models, providing mechanistic insights into how Ca^2+^ signaling regulates gap junction uncoupling and broader implications for understanding cell stress responses and tissue protection.

## INTRODUCTION

Gap junctions are intercellular communication channels, ubiquitously expressed in the human body, mediating electrical and metabolic coupling critical to tissue function and development^1,2^. Dysregulation of gap junctions contributes to a variety of diseases, including blindness, deafness, arrhythmia, stroke, and cancers^3–7^. Elevated intracellular calcium (Ca^2+^) plays a critical role in gap junction regulation by inducing channel closure, protecting healthy cells from cytotoxic signals triggered during ischemic conditions, such as heart attack or stroke^8–10^. This Ca^2+^-induced uncoupling mechanism isolates damaged cells, preventing widespread tissue damage. Despite its physiological importance, the structural and mechanistic basis of this protective gating response remains elusive.

Gap junctions exhibit a unique structural architecture that enables coordinated electrochemical communication across tissues and organs^11–14^. Intercellular channels are composed of 12 connexin subunits (21 isoforms in human^20^), each containing four transmembrane helices (TM1–4), two extracellular loops (EC1–2) for docking interactions, and an N-terminal (NT) domain involved in voltage-sensing and substrate selectivity^15–19^. Intracellular loop (ICL) and C-terminal (CT) regions involved in trafficking and regulation are largely disordered and highly variable among isoforms. Within the plasma membrane, six connexin subunits assemble into a hemichannel, which docks with a hemichannel from an adjacent cell to form a large-pore intercellular channel (∼12 Å pore diameter). This arrangement facilitates the passage of ions, metabolites, and signaling molecules^21,22^, allowing gap junction-coupled tissues to function as a syncytium, efficiently sharing long-range electrical and chemical signals across entire tissues and organs.

Elevated intracellular Ca^2+^ concentrations, reaching pathophysiological levels (*e.g.,* during ischemia, tissue injury, and apoptosis), induce closure of gap junction communication pathways independently of transjunctional voltage^23–25^. The physiological role of this Ca^2+^ gating response is distinct from hemichannels, which generally remain closed under resting membrane potentials and normal external Ca^2+^ levels (∼1.8 mM)^26,27^. However, whether the gating mechanisms underlying these channel types are fundamentally distinct remains unclear.

Early structural studies suggested Ca^2+^ induces subunit rearrangements narrowing the pore of native liver gap junctions, leading to the “iris-model” of channel gating^28–30^. This model prevailed for nearly 30 years, before being challenged by X-ray crystallography studies of connexin-26 (Cx26), which proposed an ‘electrostatic barrier’ mechanism based on the absence of large-scale structural differences observed with or without Ca^2+^ ^31^. Critically, however, these structures lack resolved NT domains, leaving the potential role of the NT in mediating Ca^2+^ gating unclear.

Functional studies on Cx26 and Cx46 hemichannels have challenged the ‘electrostatic barrier’ model, and have allosterically linked Ca^2+^ binding to the NT voltage-sensing domain^27,32–34^. However, recent cryo-EM studies of Cx31.3 hemichannels once again did not reveal any large-scale differences with or without Ca^2+^ ^35^. While the precise Ca^2+^ binding sites were unresolved in this study, the NT domains were visualized in a raised conformation that narrow the pore to ∼8 Å in both conditions. Notably, though, the lifted conformation of the NT may have been influenced by the presence of lipid or detergent molecules bound inside the pore, potentially introduced during sample preparation. Moreover, molecular dynamics simulations suggested that ion permeation could still occur^35^, confounding proposed roles for the NT as the primary gating module.

Here, we present cryo-EM structures of native lens Cx46/50 gap junction channels reconstituted in lipid nanodiscs, and exchanged to high Ca^2+^ conditions. These channels are essential for lens transparency and homeostasis^36,37^, while the formation of cataracts have been linked to calcium mishandling and aberrant Cx46/50 coupling^38–42^. Our data reveal multiple Ca^2+^ binding sites within the pore of Cx46/50 channels, inducing an ensemble of occluded and distinct gated states through NT domain conformational changes. Moreover, subunit rearrangements lead to pore collapse, facilitating steric blockade of the permeation pathway by the NT domains, reminiscent of the the original ‘iris model’. Collectively, these findings help unify decades of observations and provide critical insights into Ca^2+^-induced gating, with implications for tissue protection and connexin-related disease mechanisms.

## RESULTS

### Ca^2+^-bound occluded state of connexin-46/50

Native heteromeric/heterotypic Cx46/50 intercellular channels were purified from mammalian lens tissue and reconstituted into lipid nanodiscs containing dimyristoyl phosphatidylcholine (DMPC) and the scaffold MSP1E1 under neutral pH conditions, previously shown to result in a stabilized open-state^13,43^ (Methods). Nanodisc embedded channels were then exchanged into 20 mM Ca^2+^ buffer for further analysis, and inspected by negative stain electron microscopy to ensure homogeneity and nanodisc incorporation (Extended Data Fig. 1). These conditions were selected to ensure complete saturation of low affinity binding sites and relatively high concentrations of protein required for cryo-EM sample preparation.

Cryo-EM single particle analysis revealed Ca^2+^ binding induces a heterogenous ensemble of states. Here, we describe a Ca^2+^-bound structure of Cx46/50 representing a major class of the particles resolved to a global resolution of 2.1 Å, enabling detailed atomic modeling (Fig. 1a,b; Extended Data Fig. 2–3; Extended Data Table 1). The map revealed well-defined NT, TM1-4, and EC1/2 domains, while the intrinsically disordered CT domain and ICL regions remained largely disordered and unresolved. Bouquets of lipid chains at subunit interfaces and 100’s of ordered water molecules were also identified, parallelling features described in the 1.9 Å apo-state structure^43^.

**Figure 1:**
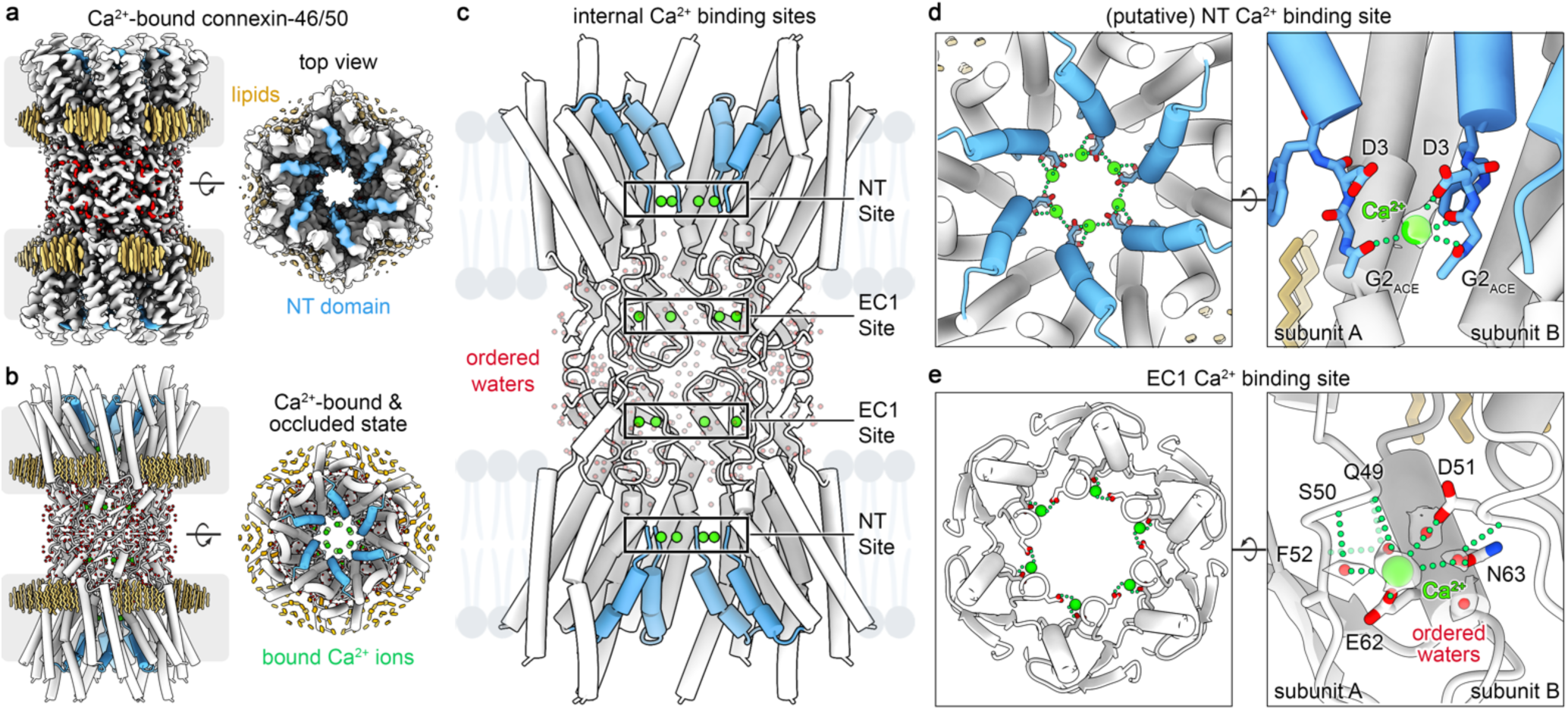
Ca2^+^-bound occluded state of connexin-46/50. **a**, Cryo-EM map and **b**, atomic model of Ca^2+^-bound occluded state of Cx46/50 gap junctions in lipid nanodiscs (protein – white; NT domain – blue; waters – red; lipids – yellow; Ca^2+^ ions – green). Gray boxes denote the approximate lipid bilayer boundaries. **c**, Internal view of the channel (lipids omitted for clarity), highlighting bound Ca^2+^ ions (green spheres) located at two binding sites within each subunit: NT and EC1 sites. **d**, Slice view (left) and zoomed view (right) of the NT site, where Ca^2+^ is coordinated by the carboxylate on D3 and the carbonyl backbone and acetylated terminus of G2_ACE_ through both intra- and inter-subunit interactions (labeled). Segmented cryo-EM density for the bound Ca^2+^ ion is shown in gray transparency. This site is putatively assigned due to the limited signal-to-noise at this site. **e**, Slice view (left) and zoomed view (right) of the EC1 site, where Ca^2+^ is coordinated by E62 and D51 on a neighboring subunit. This site is further supported by several water-mediated interactions, including with N63 and several neighboring backbone carbonyls (labeled). Segmented cryo-EM densities for Ca^2+^ and ordered waters are shown in gray transparency. Residues involved in Ca^2+^ binding are conserved in Cx46 and Cx50.

Isoform-specific assembly patterns of native heteromeric/heterotypic Cx46/50 channels could not be definitively discerned due to high sequence conservation (>80%) in structured regions, as previously described^13,43^. However, both isoforms fit equally well into the map, with sequence differences primarily localized to solvent-exposed regions (Extended Data Fig. 3–4). Accordingly, structural comparisons between resolved Ca^2+^-bound states and previously captured open apo-states use Cx46 as the primary reference, with Cx50-specific features noted where relevant.

While overall, this structure of Cx46/50 obtained under high Ca^2+^ conditions resembles the apo-state (Cα r.m.s.d. ∼0.6 Å), significant conformational changes in the NT domain are observed that partially occlude the pore entrance (Fig. 1a; Extended Data Fig. 5), further detailed in the following section. In this ‘Ca^2+^- bound occluded state’, two Ca^2+^ binding sites were identified: one at the NT domain (NT site) and another in the pore-lining extracellular loop (EC1 site) (Fig. 1c–e).

The NT site features Ca^2+^ coordination through the acetylated N-terminal G2 (G2_ACE_) and carboxylate sidechains of D3 contributed by neighboring NT domains. This arrangement forms a Ca^2+^-stabilized ring at the pore entrance, reinforcing the occluded state (Fig. 1c,d). While these features are consistent with Ca^2+^ binding, limited signal-to-noise in this region of the map leaves this assignment tentative (Extended Data Fig. 3). In contrast, the EC1 site is clearly resolved in the cryo-EM map, involving coordination by E62 and D51 carboxylate groups, further stabilized by ordered water molecules mediating interactions with N63 and EC1 backbone carbonyls (Fig. 1c,e; Extended Data Fig. 3).

Interestingly, the EC1 site is distinct from the Ca^2+^ binding site observed in Cx26^31^, highlighting potential isoform-specific differences. However, inclusion of D51 at the EC1 site aligns with prior functional mutation studies implicating this residue in Ca^2+^ binding in Cx46 and Cx26^33^. Modifying D51 would disrupt coordination with E62 and the matrix of ordered water molecules, consistent with diminished Ca^2+^ binding reported by Lopez *et al*. Notably, E62 is conserved in only four human connexins (Cx43, Cx45, Cx46, Cx50), suggesting a potentially specialized role, whereas the universal conservation of D51 implies a broader regulatory function across connexin families (Extended Data Fig. 4).

### Ca^2+^ induced changes to the permeation pathway

To further investigate the effects of Ca^2+^ binding and mechanistic implications, the permeation pathways of Cx46/50 in the open apo-state^43^ and Ca^2+^-bound occluded state were subjected to detailed structural comparison (Fig. 2). Overall, Ca^2+^ binding induces inward displacement of the distal NT domain, coupled with a clockwise rotation of the TM helices, resulting in a stabilized occluded conformation of the NT with a reduced pore aperture as well as altered electrostatic properties throughout the permeation pathway (Fig. 2a–c).

**Figure 2:**
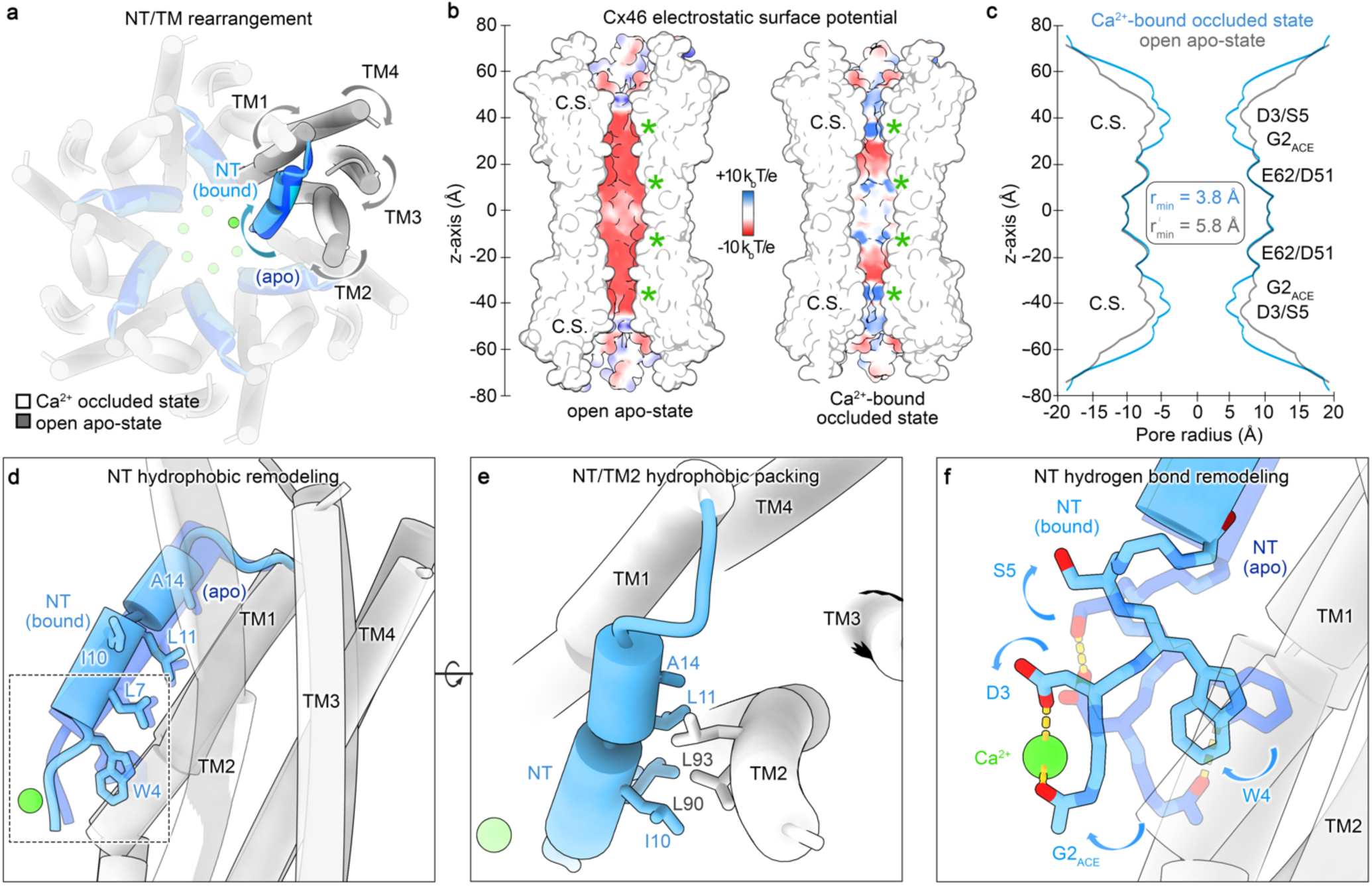
Ca2^+^ induced changes in electrostatics and conformational state of the permeation pathway. **a,** Cytoplasmic view of the channel, showing TM domain rearrangements (apo-state – grey; Ca^2+^ bound– white) and NT domain remodeling (apo-state – dark blue, Ca^2+^ bound – light blude) within each subunit. b, Coulombic surface representation of Cx46 in the open apo-state (left) and Ca^2+^-bound occluded state (right), shown in split-view to visualize the permeation pathway (positive – blue; neutral – white; negative – red). Asterisks mark Ca^2+^ binding sites, and constriction sites (C.S.) are labeled. c, Pore profile analysis of open apo-Cx46 (grey) versus Ca^2+^ bound occluded Cx46 (blue), highlighting NT domain constriction at D3/S5, respectively. Note, R9 in Cx46 (N9 in Cx50) is flexible in the open apo-state and was truncated at Cβ for this analysis. d, Conformational changes in the NT domain upon Ca^2+^ binding, with hydrophobic residues anchoring the NT to TM1/2 shown in stick representation (labeled). e, Rotated view, highlighting the stabilization of the Ca^2+^ bound NT conformation by hydrophobic interaction with TM2 involving residues L90 and L93 (V93 in Cx50). f, Zoomed view, showing remodeling of NT residues (G2 to S5) upon Ca^2+^ binding (boxed region in panel d). In the open apo-state, G2_ACE_ hydrogen bonds with W4, while D3 forms a hydrogen bond with S5. In the Ca^2+^-bound occluded state, G2_ACE_ and D3 reorient to chelate Ca^2+^, disrupting interactions with S5 and W4 and inducing their reconfiguration (arrows).

In the apo-state, the permeation pathways of both Cx46 and Cx50 are predominantly electronegative, consistent with their appreciable cation selectivity^13,18,44–47^. A band of positive charge is localized at the pore entrance in Cx46, conferred by R9 (N9 in Cx50), that has been shown to contribute to the lower ion conductance of this isoform as compared to Cx50^13,17,18^. Upon Ca^2+^ binding, the permeation pathway becomes substantially more electropositive, predictably increasing the energetic barrier for major cation permeants, such as K⁺ and Na⁺ (Fig. 2b).

The primary constriction site (C.S.) of Cx46/50 in the open apo-state is located at the proximal end of the NT domains, establishing a minimum radius (r_min_) of 5.8 Å in Cx46^48^ (Fig. 2c, grey trace; established at D3). This conformation is highly permissive to the permeation of solvated ions, supporting their relatively high conductance^13,18^. In contrast, Ca^2+^ binding induces significant conformational changes at the NT, narrowing the C.S to an r_min_ = 3.8 Å (Fig. 2c, blue trace). While this Ca^2+^-bound occluded state does not completely close the pore, such signficant narrowing would also likely contribute to altered ion conductance and potentially exclude passage to larger signaling molecules (*e.g.,* hydrated radius of cAMP ∼3.8 Å^49^).

NT remodeling associated with Ca^2+^ binding disrupts hydrophobic interactions anchoring the NT-helix to the pore vestibule (Fig. 2d). This reconfiguration is supported by a reorientation of TM2, maintaining hydrophobic contacts with residues L90 and L93 (V93 in Cx50) (Fig. 2e). Although Ca^2+^ density at the NT site is weak, conformational changes strongly support its role in Ca^2+^ coordination. In the apo-state, G2_ACE_ hydrogen bonds with the indole ring on W4, while D3 adopts an ‘upward’ orientation stabilized through hydrogen bonding with the hydroxyl group on S5^13^. Ca^2+^ binding disrupts these interactions, repositioning G2_ACE_ and D3 reorienting to a ‘downward’ state consistent with their roles in Ca^2+^ chelation. This reconfiguration is further accompanied by a pronounced reorientation of the W4 anchoring residue, consistent with a destabilization of the open-state (Fig. 2f; Supplemental Movie 1).

These structural and electrostatic changes suggest altered conductance and selectivity characteristics in the Ca^2+^-bound occluded state. However, the partially occluded ∼7.6 Å pore diameter may still permit passage of hydrated ions, implying this state may represent an intermediate configuration, rather than a fully gated state.

### Ca^2+^ binding further induces multiple gated states

Cryo-EM analysis of the entire particle dataset revealed heterogeneous NT domain density, indicating significant conformational variability. Using 3D classification methods in RELION^50^, we identified two additional Ca^2+^-bound conformational states, termed gated state 1 and gated state 2, characterized by distinct NT configurations that are fully disengaged from interactions with TM1/2 (Fig. 3a-c; Extended Data Fig. 2). In gated state 1, the proximal region of the NT domains establish a continuous ring-like interaction at the pore center, effectively sealing the permeation pathway (Fig. 3b). In gated state 2, the NT regions adopt a more lifted and kinked conformation, with proximal regions clustering deeper within the pore to form a central plug (Fig. 3c). These states likely represent fully gated conformations, effectively blocking substrate passage.

**Figure 3:**
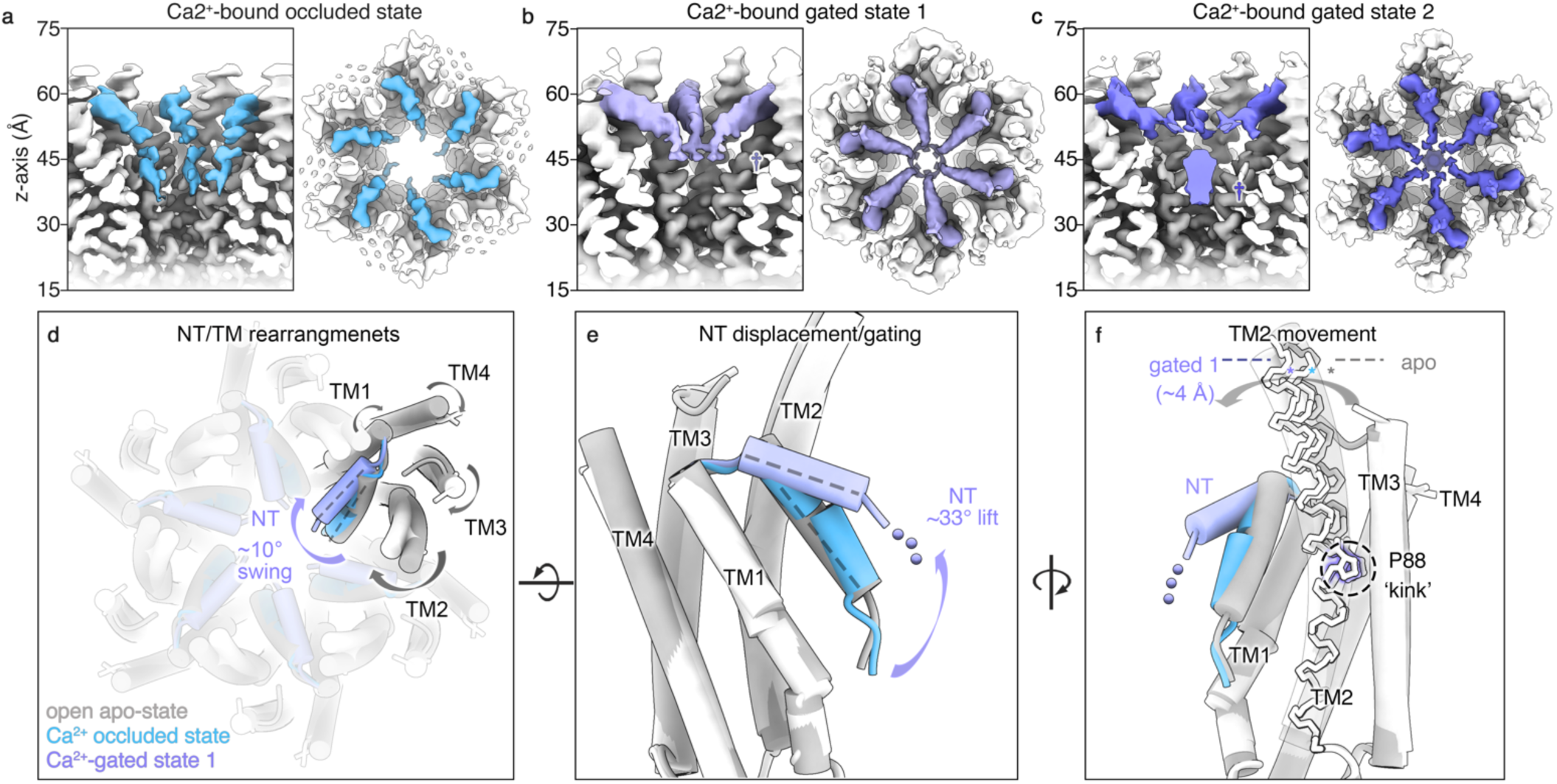
Ca2^+^ binding further induces multiple gated state conformations. **a-c,** Cryo-EM maps for the Ca^2+^-bound (a) occluded state, (b) gated state 1, and (c) gated state 2, shown in split-view. NT domains are colored blue (occluded), violet (gated state 1), and purple (gated state 2). Gated states exhibit distinct NT orientations, obstructing the pore at varying positions along the z-axis (†). **d,** Top view of the channel showing rearrangement of the TM domains that accompany movement of the NT domain (open apo state – gray, Ca^2+^-bound occluded state – light blue, Ca^2+^-bound gated state 1 – violet). Gated state 1 is distinguished by a ∼10° inward swing of the NT and reorientation of TM2 toward the pore center. **e**, Zoomed view of NT vertical displacement in gated state 1, showing a ∼33° lift along the pore axis, resulting in complete dissociation from TM1/2. f, Zoomed view of TM2 movement accompanying NT rearrangements. TM2 reorientation is facilitated by a conserved proline kink (P88), displacing the cytoplasmic end of TM2 by ∼4 Å in gated state 1 relative to the open apo-state (asterisk). TM2 undergoes similar but less pronounced movement in the Ca^2+^-bound occluded state. Unmodeled proximal NT residues in Ca^2+^-bound gated state 1 (G2 through F6) are indicated by dots in panels (e, f). An atomic model for Ca^2+^-bound gated state 2 was not built due to limited NT resolution contributing to the plug-like gate.

The refined map for Ca^2+^-bound gated state 1 (2.6 Å global resolution) supported atomic modeling up to residue 7, with sidechains for residues 7–17 truncated at the Cβ position due to limited local resolution (Fig. 3d-f; Extended Data Fig. 2-3; Extended Data Table 1). Residues 2–6 could not be confidently modeled into the ring-like feature at the center of the pore. In contrast, further refinement of Ca^2+^-bound gated state 2 failed to resolve the more substantial NT interactions with sufficient clarity to support atomic modeling.

Structural comparison of Ca^2+^-bound gated state 1 with the open apo-state revealed pronounced NT and TM domain rearrangements, similar to but more extensive than those observed in the occluded state (Fig. 3d-f). The NT undergoes a 10° swing toward the center of the pore, when viewed along the pore axis, accompanied by a 33° upward lift along the pore axis, fully detaching from the channel lumen (Fig. 3d,e; Supplemental Movie 2). These NT movements are coupled with an overall clockwise rotation of TM helices, similar to the Ca^2+^-bound occluded state, but with a more pronounced reorientation of TM2 toward the center of the pore. TM2 flexibility is facilitated by a conserved proline kink (P88 in Cx46/50), imparting flexibility to the cytoplasmic half of TM2 that enables stabilization of the various NT conformations (Fig. 3f; Extended Data Fig. 4). Additionally, minor structural variation is observed in TM1 around residues ∼39–41, which adopt a π-helix conformation. This π-helix introduces a kink in TM1, separating the para-helical region leading to EC1, and subtle variation of the hydrogen bonding at this feature compared to the open-state are observed in both the gated and occluded states (Extended Data Fig. 5).

### Ca^2+^ gating involves multiple states of NT domain closure and pore collapse

To further investigate the heterogeneity of NT conformations induced by Ca^2+^ binding, we employed 3D variability analysis (3DVA) in CryoSPARC^51^, using a symmetry-expanded particle stack. For comparison, the Cx46/50 apo-state dataset^43^ underwent the same 3DVA workflow. The apo-state structure displayed minimal conformational variability, with only slight “wobble” motions of the NT and TM2 regions, consistent with the global stability of this open-state conformation (Fig. 4a,c).

**Figure 4:**
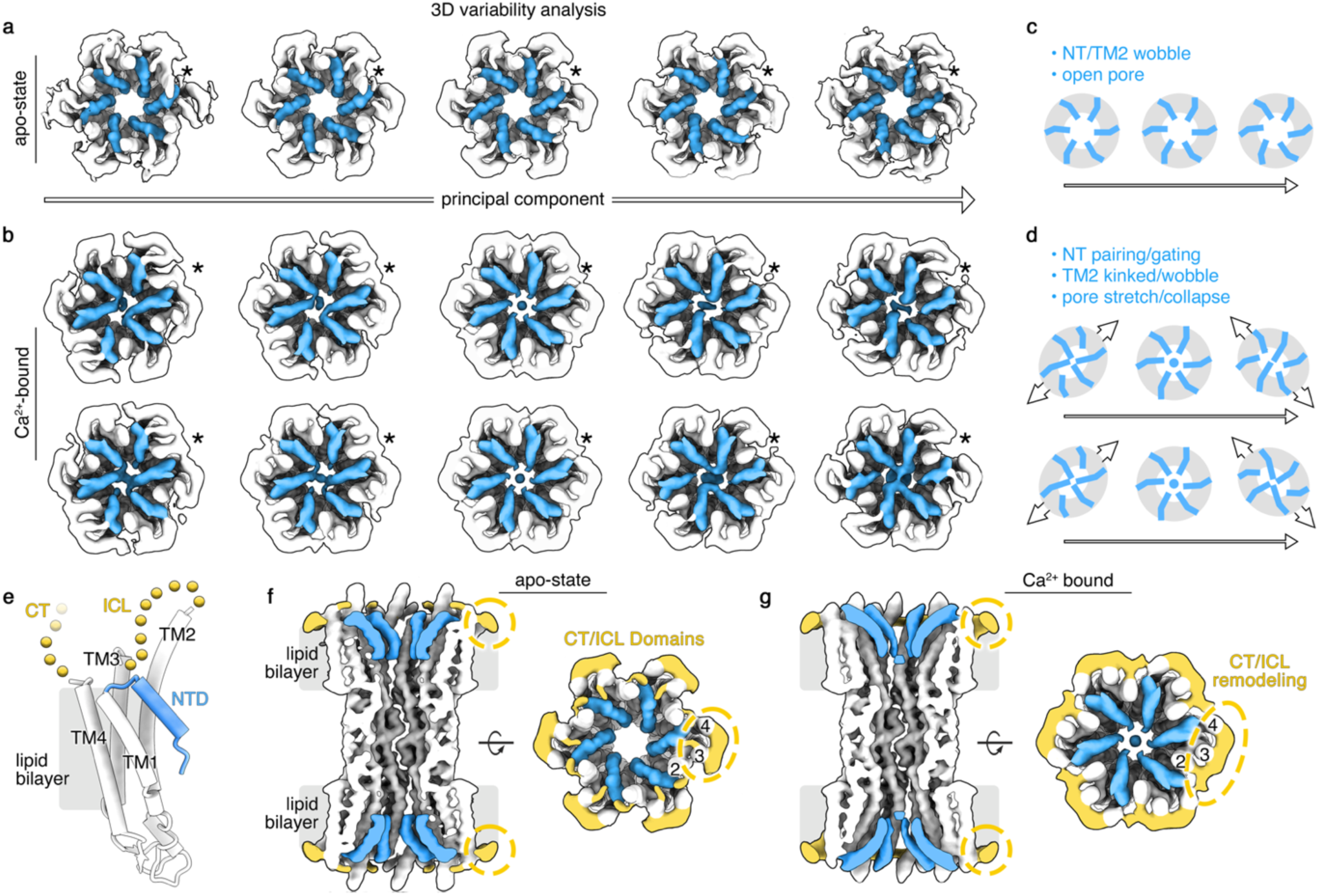
Ca2^+^-induced gating facilitated by asymmetric NT movement and pore collapse. **a-b**, Snapshots along the primary principal components (PCs) from 3D variability analysis (3DVA) for (a) the open apo-state dataset and (b) the Ca^2+^-bound dataset. NT domains are colored (blue) and TM2 for a representative subunit is highlighted (asterisk). **c-d**, Summary and schematics of observed domain movements. The open apo-state dataset shows minimal variability, with NT and TM2 domains exhibiting a slight “wobble” and a stable open pore. In the Ca^2+^-bound dataset, NT domains show significant rearrangements, transitioning between occluded and gated states that block the pore. Gated states involve various NT pairing interactions, either between neighboring or opposing subunits stretching across the channel pore. These interactions are facilitated by pore stretching and collapse, which reduce the cross-pore distance, facilitating NT interactions. **e**, Model of a single Cx46/50 subunit, illustrating unmodeled intracellular loop (ICL) and C-terminal (CT) domains (yellow dots). **f-g**, Representative cryo-EM maps from the (f) apo-state dataset and (g) Ca^2+^-bound dataset taken from the center of the PCs. Map densities corresponding to unmodeled ICL/CT regions (yellow) show minimal variability in the apo-state dataset (compare panel f and a) but undergo significant reorganization along the 3DVA PCs in the Ca^2+^ bound dataset (compare panel g and b), indicating coupled conformational changes with gated states.

In contrast, 3DVA of the Ca^2+^-bound dataset revealed continuous NT domain dynamics, delineated by multiple principal components, including clear deviations from the D6 symmetry characteristic of these dodecameric channels (Fig. 4b; Extended Data Fig. 6; Supplemental Movie 3). At the extremes of these principal components, a variety of Ca^2+^-bound conformational states emerged, characterized by some general principles. Notably, only a subset of NT domains adopted a gated state conformation, while others remained in an occluded (or potentially open) state. Gated NT domains exhibited paired interactions, typically involving 2–4 subunits. NT-pairing interactions occur either laterally between neighboring subunits or across the channel between opposing subunits, effectively blocking the pore (Fig. 4b). Traversing the principal components, the NT domains transition to symmetrical configurations resembling the Ca^2+^-bound occluded state (compare Fig. 4b, center to Fig. 3a).

These NT movements were accompanied by subunit reconfigurations that result in an overall stretching of the channel framework, leading to pore collapse along the orthogonal axis. This structural remodeling facilitates NT interactions between opposing subunits, culminating in an obstructed permeation pathway (Fig. 4b,d; Extended Data Fig. 6). Additionally, unmodeled regions of the cryo-EM maps corresponding to regions of the intracellular loop (ICL) and/or C-terminal (CT) domains displayed conformational changes correlated with NT dynamics, suggesting coupled structural rearrangements (Fig. 4e,g; Extended Data Fig. 6; Supplemental Movie 3). By contrast, no significant NT dynamics or pore collapse was observed by 3DVA of the apo-state, and the ICL/CT regions exhibited minimal variability (Fig. 4a,b,f).

Together, these findings indicate that Ca^2+^ binding drives collective conformational dynamics that play an integral role in the mechanism of channel inhibition, encompassing multiple states that include NT-domain closure and pore collapse. The diverse array of gated states uncovered reveal the underlying complexity of the Ca^2+^-induced mechanism, involving a broad spectrum of dynamic NT configurations rather than a simple two-state gating process.

## DISCUSSION

### Ca^2+^ induced gating in Cx46/50 gap junctions

Our structural analysis demonstrates that Ca^2+^ binding at multiple sites induces conformational changes in Cx46/50 gap junctions, particularly within the NT domains, leading to pore occlusion and closure (Fig. 5). In the absence of Ca^2+^, or other gating stimuli, Cx46/50 adopts a stabilized open-state conformation, where the NT forms an amphipathic α-helix anchored to the channel lumen by hydrophobic interactions with TM1/2. These stabilizing interactions maintain a large-pore permeation pathway (∼12 Å diameter), consistent with electrophysiological data^18^. Ca^2+^ binding disrupts these stabilizing interactions, driving NT remodeling and transitioning the channel into an ensemble of occluded and gated states.

**Figure 5:**
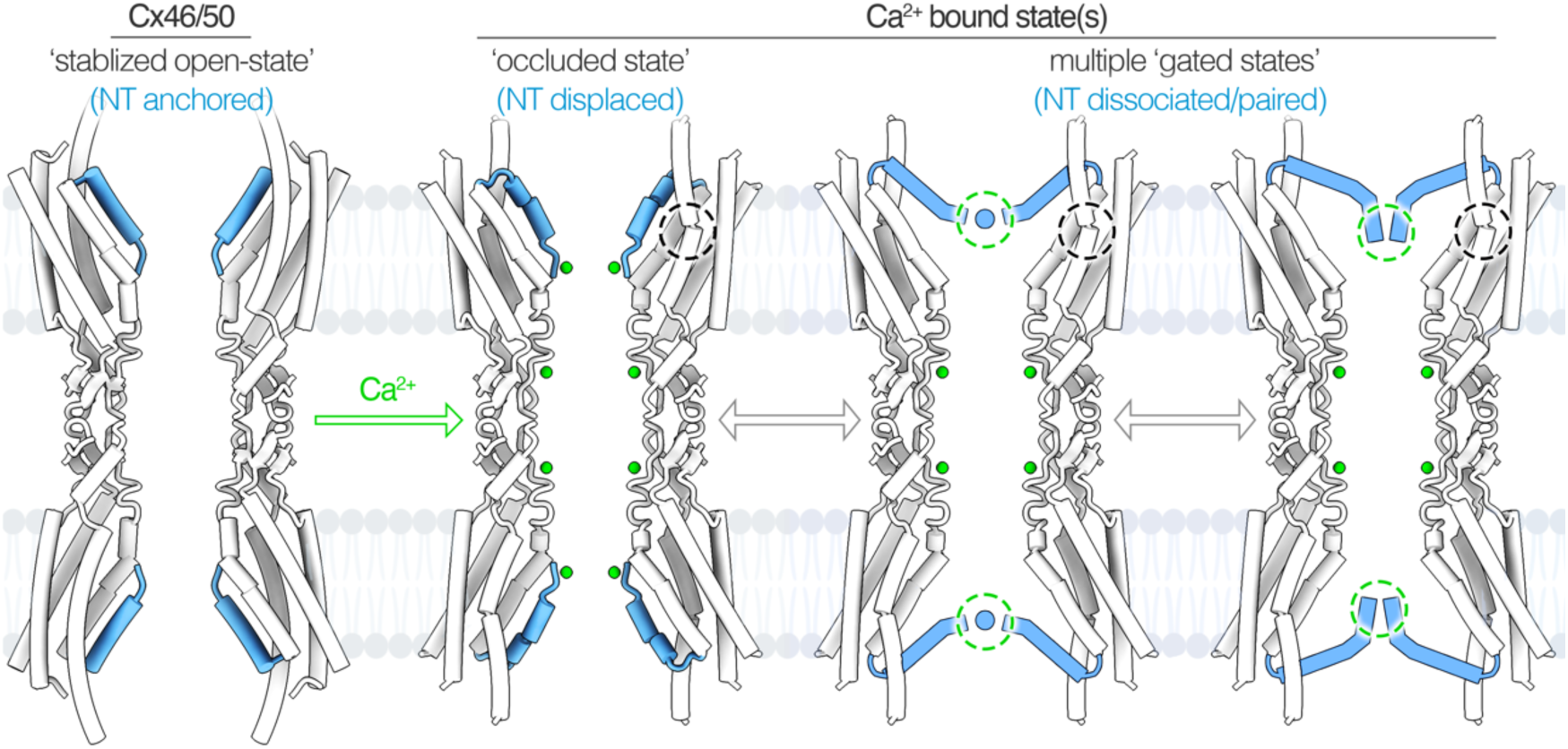
Overview of Ca^2+^-induced pore occlusion and gating in connexin-46/50 gap junctions. In the apo-state, Cx46/50 channels adopt a stabilized open-state, with NT domains (blue) anchored to the channel lumen via hydrophobic interactions with TM1/2. Under high Ca^2+^ conditions, Ca^2+^ ions (green) bind to two sites per subunit: a putative site at the NT domain and a well-defined site on the EC1 domain. Ca^2+^ binding induces NT conformational changes supported by TM2 reorientation, facilitated by a conserved proline kink (dashed black circles). These structural rearrangements result in an ensemble of occluded and gated states, with altered pore electrostatics, where NT domains are proposed to reduce or block ion and small molecule permeation. NT domain pairing contributes to steric blockade of the permeation pathway, facilitated by subunit rearrangements that collapse the pore. It is proposed that NT paring may also be supported by inter-subunit Ca^2+^ binding (dashed green circles).

In the Ca^2+^-bound occluded state, Ca^2+^ binding at the NT site is supported by interactions with the acetylated G2 and conserved D3 residues. While the NT remains engaged with TM1/2, remodeling of hydrophobic anchoring residues leads to its displacement and constriction of the pore. The additionally formed gated states feature complete NT disengagement from TM1/2 and pore blockage via proximal NT interactions. Synergistic reorientations of TM2, facilitated by a tightly conserved proline (P88 in Cx46/50), further stabilize these various NT conformations (Fig. 5, black circle). Proline positions within transmebrane helices play functional roles in facilitating signal transduction in many ion channels and receptors^52^. Notably, a homologous proline site has been assoicated with modulating the voltage-gating properties of Cx26 and Cx32^53,54^, indicating a potentially conserved role of both the NT and TM2 in transducing gap junctional gating in response to diverse physiological signals.

Detailed heterogeneity analysis highlighted the dynamic nature of Ca^2+^-induced gating, revealing a spectrum of NT configurations, featuring paired NT interactions that contribute to the ensemble of gated states. This analysis further uncovered dynamic modes of channel stretching enabled by subunit rearrangements that culminate in pore collapse, facilitating NT-pairing interactions across the channel pore. The ensemble of gated states underscores the complexity of the Ca^2+^-induced mechanism, extending beyond a simple binary open-closed model to encompass a dynamic array of NT configurations. This observation is consistent with the graded Ca^2+^-induced closure to larger molecules versus small ions^55^, or formation of sub-conductance states^18,56^. Additionally, coupled rearrangments of the ICL and/or CT domains suggest a broader structural network of interactions involved in channel gating, expanding the mechanistic framework for regulation beyond the NT and well-ordered TM/EC domains.

The provisional NT Ca^2+^ binding site aligns with observed rearrangements of the proximal NT region (G2–W4) in the Ca^2+^-bound occluded state, resulting in destabilization of hydrophobic anchoring with TM1/2. Intriguingly, shared Ca^2+^ binding at the NT site could also plausibly explain NT-pairing in the gated states through sub-stoichiometric Ca^2+^ coordination, where a single Ca^2+^ ion could effectively cross-link neighboring or opposing NT domains (Fig. 5, green circles). Shared Ca^2+^ coordination could effectively drive displacement of the NT hydrophobic anchoring residues from the channel lumen, overcoming the energetic costs of exposing these hydrophobic regions to solvent.

Under physiological Ca^2+^ concentrations that result in channel gating in cells (high nanomolar to micromolar), these sub-stoichiometric gated states may dominate in gap junctional uncoupling. However, the ambiguitiy of the NT Ca^2+^ binding and proposed role in facilitating NT-paring warrant further experimental validation. Of note, the functional role of D3 in channel conductance and voltage-sensing^16,57–59^, and the dependence of N-terminal acetylation on the residue type at position 2^60^ make targeted mutational studies challenging, as alteration of these sites may inadvertently disrupt other critical aspects of connexin channel function.

### Insights into proposed Ca^2+^ gating models

Our findings reconcile key elements of proposed Ca^2+^ gating models, resolve long-standing ambiguities, and point to directions for future investigation. While the EC1 Ca^2+^ binding site in Cx46/50 differs from the location resolved in the Ca^2+^-bound Cx26 structure^31^, both induce alterations in pore electrostatics that would impede cation permeation. The pronounced NT flexibility observed in Cx46/50 also provides a plausible explanation for the unresolved NT domains in the Cx26 study, where structural heterogeneity likely obscured this critical gating element. Early electron crystallographic studies^15,61^ and recent investigations into the structural mechanisms of pH and CO_2_ regulation of Cx26 further support the role of the NT as a dynmaic gating module^62–64^. Structural variability of the NT has also been documented for Cx43, Cx32 and Cx36, highlighting its dynamic nature across connexin families^65^.

Functional studies on Cx46 hemichanels suggest Ca^2+^ binding is allosterically coupled to the voltage-sensing mechanism (thought to be mediated by the NT, and specifically involving D3^19,58^), resulting in a physical constriction of the pore^34^. The pivotal role of the NT domain in Ca^2+^-gating is further supported by pore-accessibility studies conducted on Cx26 and Cx46 hemichannels, which demonstrate that the EC1 region of the pore remains accessible in the presence of Ca^2+^^33^. This work also implicates D51 in Ca^2+^ binding for Cx46, validating the interactions resolved at the EC1 Ca²⁺ binding site in our study. D51 (D50 in Cx26) is highly conserved across most human connexins, and mutations such as D50N in Cx26, linked to keratitis-ichthyosis-deafness (KID) syndrome, result in loss of hemichannel Ca^2+^ regulation, further supporting its functional significance^32,66^.

The role of E62 in Ca^2+^ binding for Cx46/50, as resolved in this study, aligns with prior molecular dynamics simulations suggesting that this site forms quasi-stable interactions with monovalent cations^67^. However, the minimal conservation of a negatively charged residue at this site suggests it may contribute to isoform-specific features of Ca^2+^ sensitivity. Likewise, sequence diversity at key positions such as D50 and D3 among a few other Ca^2+^-sensitive isoforms, such as Cx32 and Cx36, further highlights the likelihood for isoform-specific gating properties. Furthermore, the role of auxiliary Ca^2+^-sensing proteins, such as calmodulin, may add additional layers of regulation, contributing to the diversity of Ca^2+^-mediated modulation of connexin channel function^68^.

Overall, this work underscores the key role played by the NT domain and the complexity of underlying dynamics involved in Ca^2+^ gating of Cx46/50. Remarkably, the Ca^2+^-induced pore collapse observed in our study appears to be a key feature facilitating the NT gating mechanism, evoking elements of the “iris model” of channel gating originally proposed by Zampighi, Unwin, and Ennis over 40 years ago^28–30^. Together, these findings contribute to a more unified framework for understanding Ca^2+^ gating in connexin gap junctions, highlighting an intricate interplay of isoform-specific adaptations and conserved structural mechanisms.

### Broader implications

Gaining a clear mechanistic picture of the Ca^2+^ gating response in the gap junctions is critical to understanding how tissues are protected from localized trauma or stress. Without dedicated mechanisms to uncouple damaged cells, cytotoxic signals could propagate through gap junctions, triggering widespread cell death—a phenomenon aptly termed the “bystander effect” or “kiss of death“^69,70^. This protective mechanism is thought to play a vital role in minimizing tissue damage during conditions of calcium overload associated with heart attack and stroke. Conversely, within the eye lens Ca^2+^-induced closure of Cx46/50 would be an aberrant consequence of aging, leading to cascading effects of cataract formation. Our findings therefore illuminate the structural underpinnings of Ca²⁺ gating, but also underscore the therapeutic potential of targeting connexin gating mechanisms^7^. Developing interventions to modulate gap junction uncoupling may offer promising strategies for treating connexin-linked conditions, including heart disease, stroke, and cataract.

## ACKNOWLEDGEMENTS

We thank Dr. Lisa Ebihara and Dr. Bassam Haddad for helpful discussions. We are grateful for instrumentation access and training provided by the staff at the OHSU Multiscale Microscopy Core and Advanced Computing Center, and the Pacific Northwest Cryo-EM Center (supported by NIH Grant R24GM154185) with support from Dr. Janette Myers. The research was funded by NIH grants R35GM124779 (to S.L.R.) and fellowship F31EY030409 (to J.A.F.).

## AUTHOR CONTRIBUTIONS

J.A.F. and S.E.O. contributed equally to the work. J.A.F., S.E.O., and S.L.R. contributed to the conception and experimental design of the work. S.E.O. conducted the protein purification, nanodisc reconstitution, preparation of cryo-EM specimens and collected the cryo-EM datasets. S.E.O. and J.A.F. performed image analysis. J.A.F. performed cryo-EM classification and variability analysis. J.A.F. and J.M.J. performed atomic modeling. J.A.F., S.E.O., and J.M.J. contributed to structural interpretation. J.A.F. and S.L.R. wrote the first draft of the paper, and all authors contributed to revisions of the manuscript.

## CONFLICT OF INTERESTS

Authors declare no competing interests.

## METHODS

### MSP expression and purification

The MSP1E1 expression plasmid was obtained from Addgene^71^ and the protein was expressed and purified as previously described^43^. Freshly transformed *E. coli* cells (BL21Gold-DE3) were cultured in LB medium containing 50 μg mL^-1^ kanamycin at 37°C with shaking (250 rpm). Protein expression was induced with 0.5 mM Isopropyl β-d-1-thiogalactopyranoside (IPTG) at an OD_600_ of ∼0.5–0.6, and allowed to proceed for 3–5 hours post-induction at 37°C. Cells were harvested by centrifugation at 4,000 × *g* for 20 minutes at 4°C, and the resulting cell pellets were resuspended in MSP Lysis Buffer (40 mM Tris [pH 7.4], 1% Triton X-100, 1 mM PMSF) at ∼20 mL buffer per liter of culture. The resuspended cells were flash-frozen in liquid nitrogen and stored at -86°C for later use.

Frozen cell suspensions were thawed, supplemented with 1 mM phenylmethylsulfonyl fluoride (PMSF), and lysed by sonication on ice. The crude lysate was clarified by ultracentrifugation at 146,550 × *g* for 30 minutes at 4°C. The supernatant was filtered through a 0.22 μm membrane (Millipore) and loaded onto a gravity column containing 5 mL of HisPur Ni-NTA resin (Thermo Fisher Scientific) pre-equilibrated with Equilibration Buffer (40 mM Tris [pH 7.4]). The resin was sequentially washed with 5 column volumes (CV) of each of the following buffers: Equilibration Buffer, Triton Buffer (40 mM Tris [pH 8.0], 300 mM NaCl, 1% Triton X-100), Cholate Buffer (40 mM Tris [pH 8.0], 300 mM NaCl, 50 mM cholate), and Imidazole Wash Buffer (40 mM Tris [pH 8.0], 300 mM NaCl, 50 mM imidazole). MSP1E1 was eluted with 3 CV of Elution Buffer (40 mM Tris [pH 8.0], 300 mM NaCl, 750 mM imidazole). The eluate was filtered (0.22 μm, Millipore) and subjected to gel filtration chromatography on a BioRad ENC70 column equilibrated in 20 mM HEPES (pH 7.4), 150 mM NaCl, and 1 mM EDTA using an FPLC system (BioRad NGC). Peak fractions were identified by UV absorbance at 280 nm, pooled, and concentrated to 400– 600 μM using centrifugal concentrators. Final protein concentration was determined by UV_280_, and samples were aliquoted, flash-frozen in liquid nitrogen, and stored at -86°C for long-term use.

### Cx46/50 purification and nanodisc reconstitution

Native Cx46/50 intercellular channels were isolated from ovine lens fiber cells^13^. Fresh lamb eyes obtained from Wolverine Packers slaughterhouse (Detroit, MI) were dissected and intact lenses were stored at -86°C until use. Gap junctions were purified from lens core fiber cell tissue, which are enriched in the C-terminal truncation variant of Cx46/50 (MP38)^72–77^. Details of the purification procedure are provided below.

Lenses were thawed from -86°C, and core tissue was dissected from the cortex using a surgical blade. Stripped membranes were prepared following established protocols^78–80^. Protein concentration was measured using the BCA assay (Pierce), and membranes were stored at -86°C in storage buffer (10 mM Tris [pH 8.0], 2 mM EDTA, 2 mM EGTA) at ∼2 mg mL^-1^.

For Cx46/50 purification, stripped membranes were solubilized in storage buffer containing 20 mg mL^-1^ n-decyl-b-D-maltoside (1% wt vol^-1^) (DM; Anatrace) at 37°C for 30 minutes with gentle agitation. Insoluble material was removed by ultracentrifugation (∼150,000 × *g*, 30 minutes, 4°C), and the filtered supernatant (0.22 μm; Millipore) was subjected to ion-exchange chromatography (UnoQ; BioRad). Bound protein was eluted using a 25 CV gradient from Buffer A (10 mM Tris [pH 8.0], 2 mM EDTA, 2 mM EGTA, and 0.3% DM wt vol^-1^) to Buffer B (Buffer A + 500 mM NaCl). Peak fractions containing Cx46/50, verified by SDS-PAGE, were pooled and subjected to gel filtration chromatography on a Superose 6 Increase 10/300 GL column (GE Healthcare) equilibrated in GEL FILTRATION buffer (20 mM HEPES [pH 7.4], 150 mM NaCl, 2 mM EDTA, 2 mM EGTA, and 0.3% DM wt vol^-1^). Peak fractions were concentrated to ∼5 mg mL^-1^ using a 50 kDa m.w.c.o. centrifugal device (Vivaspin 6; Sartorius) and quantified by UV_280_ absorbance.

Freshly purified Cx46/50 was reconstituted into MSP1E1 nanodiscs with dimyristoyl phosphatidylcholine (DMPC; Avanti) following established procedures^43,81,82^. DMPC in chloroform was dried under N_2_ gas, then placed under vacuum overnight to remove residual solvent. The resulting thin film was resuspended in 50 mg mL^-1^ DM and sonicated at 37°C for 2 hours. Freshly purified Cx46/50 and DMPC were combined at a 0.6:90 (protein:lipid) molar ratio. The mixture was incubated at 25°C with gentle agitation for 2 hours. MSP1E1 was then added to achieve a final molar ratio of 0.6:1:90 (Cx46/50:MSP1E1:lipids), and the solution was incubated at 25°C for 30 minutes. Detergent was removed using SM-2 Bio-Beads (BioRad) added at a 20:1 beads:detergent (wt wt^-1^) ratio by overnight incubation (∼16 hours) at 25°C with gentle agitation. Bio-Beads were removed by perforating the top and bottom of the Eppendorf tube with a hot needle and gently centrifuging (∼500 × *g*) into a new tube containing fresh Bio-Beads (20:1 wt/wt). The second incubation was performed for an additional 2 hours at 25°C.

After Bio-Bead incubation, the samples were ultracentrifuged at ∼150,000 × *g* for 15 minutes at 4°C to remove insoluble material. The supernatant was filtered (0.22 μm; Millipore) and subjected to gel filtration chromatography using a Superose 6 Increase 10/300 GL column (GE Healthcare). GEL FILTRATION was performed in detergent-free buffer containing 20 mM Ca²⁺ to exchange Cx46/50-nanodiscs into a high-Ca²⁺ environment and to remove empty nanodiscs. Peak fractions containing Cx46/50 incorporated into nanodiscs, confirmed by SDS-PAGE, were pooled and concentrated to ∼2.5 mg mL^-1^ using a 50-kDa cut-off centrifugal filter (Vivaspin 6; Sartorius). Protein concentration was determined by UV absorbance at 280 nm. All chromatography steps were performed using FPLC at 4°C.

### Cryo-EM specimen preparation and data collection

Cx46/50-nanodiscs in 20 mM Ca^2+^ were prepared for cryoEM by applying 5 μl of sample (∼2.1 mg mL^-1^) to a glow-discharged holey carbon grid (Quantifoil R 2/1, 400 mesh) at 100% humidity. After an 8.0-second wait time, grids were blotted for 5.0 seconds, followed by a 3.0-second dwell time, and plunge-frozen into liquid ethane using a Vitrobot Mark IV (Thermo Fisher Scientific). Frozen grids were stored under liquid nitrogen until imaging.

Cryo-EM imaging was conducted on a Titan Krios (Thermo Fisher Scientific) operated at 300 kV. Dose-fractionated image stacks were acquired using a K3 direct electron detector (Thermo Fisher Scientific) in super-resolution mode, with a nominal magnification of 120,000x, corresponding to a physical pixel size of 0.830 Å (0.415 Å in super-resolution). Images were acquired at nominal dose rate of 0.49 e^-^ Å^-2^ sec^-1^, with a total dose of ∼37 e^-^ Å^-2^. A total of 5,750 movies were collected at defocus values ranging from ∼0.5–1.5 μm. Data collection was performed in an automated fashion using SerialEM^83^.

### Cryo-EM image processing for Cx46/50-lipid nanodiscs in 20 mM Ca^2+^

Beam-induced motion correction and contrast transfer function (CTF) estimation were performed in CryoSPARC v4.2.1 (Structura Biotechnology)^84,85^. Micrographs with CTF models worse than 5 Å resolution were discarded, leaving 5,205 micrographs for further processing. Initial particle picking via CryoSPARC’s blob picker yielded 3,940,202 particles, which were subjected to 2D classification to produce a cleaned stack of 592,261 particles. These particles were subjected to multiclass ab initio reconstruction with four classes, followed by non-uniform refinement with D6 symmetry, producing a 2.2 Å map from three top classes (306,198 particles, respectively; 256-pixel box, 1.038 Å pixel^-1^).

Forty projections from the non-uniform refinement map were used for template-based particle picking, generating 3,890,521 picks. Four rounds of 2D classification reduced the dataset to 1,043,250 true particles. Subsequent heterogeneous refinement and duplicate particle removal (100 Å minimum separation) yielded a dataset of 675,531 particles (280-pixel box, 0.947 Å pixel^-1^), which refined to a 2.08 Å resolution map after non-uniform refinement with D6 symmetry. The refined stack was converted to RELION^86^ format using UCSF PyEM (v0.5)^87^ for further processing. Symmetry expansion of a randomized subset of particles, yielding a 1,200,000 expanded particle set, was also prepared for 3D variability analysis (3DVA) in CryoSPARC^51^.

In RELION v4.0^88^, beam-induced motion correction, CTF estimation, and 3D auto-refine with D6 symmetry yielded a 2.55 Å map (240-pixel box, 0.968 Å pixel^-1^) on the refined stack of particles. An initial atomic model was fit into the unsharpened map, and an 8 Å-resolution map simulated from the model using UCSF ChimeraX^89^ was used to derive a solvent mask for 3D classification without alignment. This classification identified three distinct Ca^2+^-bound conformational states (occluded, gated 1, and gated 2), distinguished by NT domain features. Additional classes with apprently mixed NT conformations or missing NT density were also observed (Extended Data Fig. 2). Exploration of various symmetries or asymmetric refinements did not improve map resolution for these classes.

Bayesian polishing and CTF refinement (including per-particle defocus, beam-tilt, astigmatism, and higher-order aberration corrections) followed by 3D auto-refinement were applied to pooled particles for the occluded, gated 1, and gated 2 states. The Ca^2+^-bound occluded state (242,797 particles) refined to 2.2 Å resolution (gold-standard FSC). The Ca^2+^-bound gated 1 state (173,079 particles) resolved to 2.6 Å, and the gated 2 state (89,156 particles) refined to 2.9 Å. Postprocessing and local resolution estimation were performed in RELION, with local resolution-filtered maps generated for atomic model refinement. A full summary of the image processing workflow and resolution assessments is provided in Extended Data Figs. 2,3.

### Atomic modelling, refinement, and validation

Previously determined atomic models of Cx46 (PDB: 7JKC) and Cx50 (PDB: 7JJP) in the apo-state^43^ were rigid-body fit into maps for the Ca^2+^-bound occluded and gated 1 states. The gated 2 state was not modeled due to insufficient resolution of individual NT gating domains. Lipid acyl chains and solvent molecules were removed from the initial models, and all-atom models for Cx46 and Cx50 underwent iterative manual adjustments in COOT^90^ and real-space refinement in PHENIX^91^. Secondary structure and non-crystallographic symmetry (D6) restraints were applied during refinement, and model quality was assessed after each iteration using MolProbity^92^. Coordinate and restraint files for DMPC (PDBe Ligand Code: MC3) were generated in PHENIX eLBOW^93^. DMPC ligands were manually fit into cryo-EM density maps using ChimeraX and COOT, and unresolved portions of the ligands were deleted. Refinement was iteratively performed on the entire model until convergence of refinement statistics was achieved (Extended Data Table 1).

### Sequence and structural comparisons

Primary sequence alignments were performed using Clustal-w^94^ and viusalized in Jalview^95^. Structural alignments and analysis of structural and electrostatic properties were performed in ChimeraX^89^. Pore profile analysis was performed using HOLE^48^. For this analysis, the sidechain of R9 on Cx46 (7JKC) was pruned to Cβ to account for the dynamic nature of this resdiue as demonsrated by molecular dynamics simulation^18^ and the weak density of the sidechain in the original cryo-EM map^13,43^. This generated a primary constriction site at residue D3, consistent with Cx50 (7JJP) and (ref: 18).

### Figure preperation

Structural models and cryo-EM density maps were visualized and prepared for presentation using ChimeraX^89^. Final figures were composed in Photoshop.

### AI-assisted technologies

During the preparation of this work the authors used ChatGPT to help revise portions of the text to improve readability. After using this tool, the authors reviewed and edited the content as needed and take full responsibility for the content of the publication.

## DATA AVAILABILITY

Cryo-EM density maps have been deposited to the Electron Microscopy Data Bank (EMD-XXXX: Cx46 Ca^2+^ occluded; EMD-XXXX: Cx50 Ca^2+^ occluded; EMD-XXXX: Cx46 Ca^2+^ gated; EMD-XXXX: Cx50 Ca^2+^ gated). Coordinates for atomic models have been deposited to the Protein Data Bank (PDB: XXXX: Cx46 Ca^2+^ occluded; PDB: XXXX: Cx50 Ca^2+^ occluded; PDB: XXXX: Cx46 Ca^2+^ gated; PDB: XXXX: Cx50 Ca^2+^ gated). The original multi-frame micrographs have been deposited to EMPIAR (EMPIAR-XXXXX). Previously published models of Cx46 and Cx50 used for comparative analysis and intial modeling can be found here: (PDB 7JKC) and (PDB 7JJP).

## EXTENDED DATA TABLES AND FIGURES

**Extended Data Table 1:** Cryo-EM data collection, refinement, and validation statistics for the Cx46/50 Ca^2+^-bound dataset.

| State:<br>Isoform: | Ca<sup>2+</sup> occluded<br>Cx46 | Ca<sup>2+</sup> Occluded<br>Cx50 | Ca<sup>2+</sup> Gated 1<br>Cx46 | Ca<sup>2+</sup> Gated 1<br>Cx50 |
| --- | --- | --- | --- | --- |
| **Data collection and processing** |  |  |  |  |
| Magnification | 120,000x | 120,000x | 120,000x | 120,000x |
| Voltage (kV) | 300 | 300 | 300 | 300 |
| Electron Exposure (e-/Å<sup>2</sup>) | 37 | 37 | 37 | 37 |
| Defocus Range (μm) | -0.5 to -1.5 | -0.5 to -1.5 | -0.5 to -1.5 | -0.5 to -1.5 |
| Pixel size (Å) | 0.415 | 0.415 | 0.415 | 0.415 |
| Symmetry imposed | D6 | D6 | D6 | D6 |
| Initial particle images (no.) | 669,045 | 669,045 | 669,045 | 669,045 |
| Final particle images (no.) | 242,797 | 242,797 | 173,074 | 173,074 |
| Map resolution (Å) | 2.2 | 2.2 | 2.6 | 2.6 |
| FSC threshold | 0.143 | 0.143 | 0.143 | 0.143 |
| Map resolution range (Å) | 2.1 - 4.0 | 2.1 - 4.0 | 2.4 - 4.4 | 2.4 - 4.4 |
| **Refinement** |  |  |  |  |
| Initial model used (PDB code) | 7JKC | 7JJP | 7JKC | 7JJP |
| Model resolution (Å) | 2.3 | 2.2 | 2.7 | 2.7 |
| FSC threshold | 0.5 | 0.5 | 0.5 | 0.5 |
| Model resolution range (Å) | 2.3 - 2.4 | 2.2 - 2.3 | 2.7 - 2.8 | 2.7 - 2.8 |
| Map sharpening *B*-factor (Å<sup>2</sup>) | -42.65 | -42.65 | -32.27 | -32.27 |
| Model composition |  |  |  |  |
| Non-hydrogen atoms | 20,820 | 20,784 | 19,188 | 19,008 |
| Protein residues | 2280 | 2280 | 2244 | 2220 |
| Ligands | 192 | 192 | 156 | 156 |
| *B* factors (Å<sup>2</sup>) |  |  |  |  |
| Protein | 30.00 | 38.82 | 53.75 | 61.16 |
| Ligand | 68.55 | 68.55 | 68.55 | 68.55 |
| R.M.S. deviations |  |  |  |  |
| Bond lengths (Å) | 0.009 | 0.007 | 0.002 | 0.003 |
| Bond angles (°) | 0.705 | 0.783 | 0.472 | 0.708 |
| Validation |  |  |  |  |
| MolProbity score | 1.18 | 1.16 | 1.21 | 1.07 |
| Clashscore | 3.96 | 3.74 | 2.79 | 2.18 |
| Poor rotamers (%) | 0.00 | 0.58 | 0.00 | 1.26 |
| Ramachandran plot |  |  |  |  |
| Favored (%) | 98.38 | 98.38 | 97.27 | 98.90 |
| Allowed (%) | 1.62 | 1.62 | 2.73 | 1.10 |
| Disallowed (%) | 0.0 | 0.0 | 0.0 | 0.0 |

**Extended Data Figure 1:**
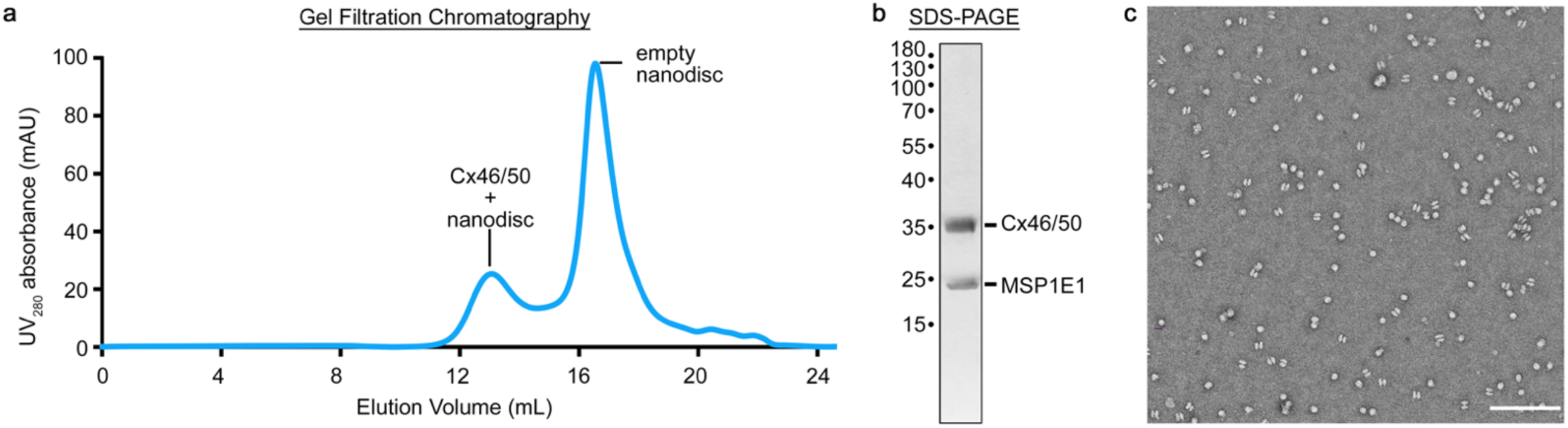
Cx46/50 reconstitution into MSP1E1/DMPC lipid nanodiscs with 20 mM Ca^2+^. **a**, Gel filtration chromatography trace (UV_280_) showing peak fractions corresponding to Cx46/50 reconstituted into MSP1E1 nanodiscs and empty nanodiscs. **b**, Silver stained SDS-PAGE of peak fractions confirms MSP1E1 (∼25 kDa) and co-migrating Cx46/50 (∼35 kDa) proteins, reflecting natively truncated C-terminal forms from lens fiber cells. **c**, Representative electron micrograph of negatively stained Cx46/50 reconstituted in MSP1E1 nanodiscs. Scale bar = 100 nm.

**Extended Data Figure 2:**
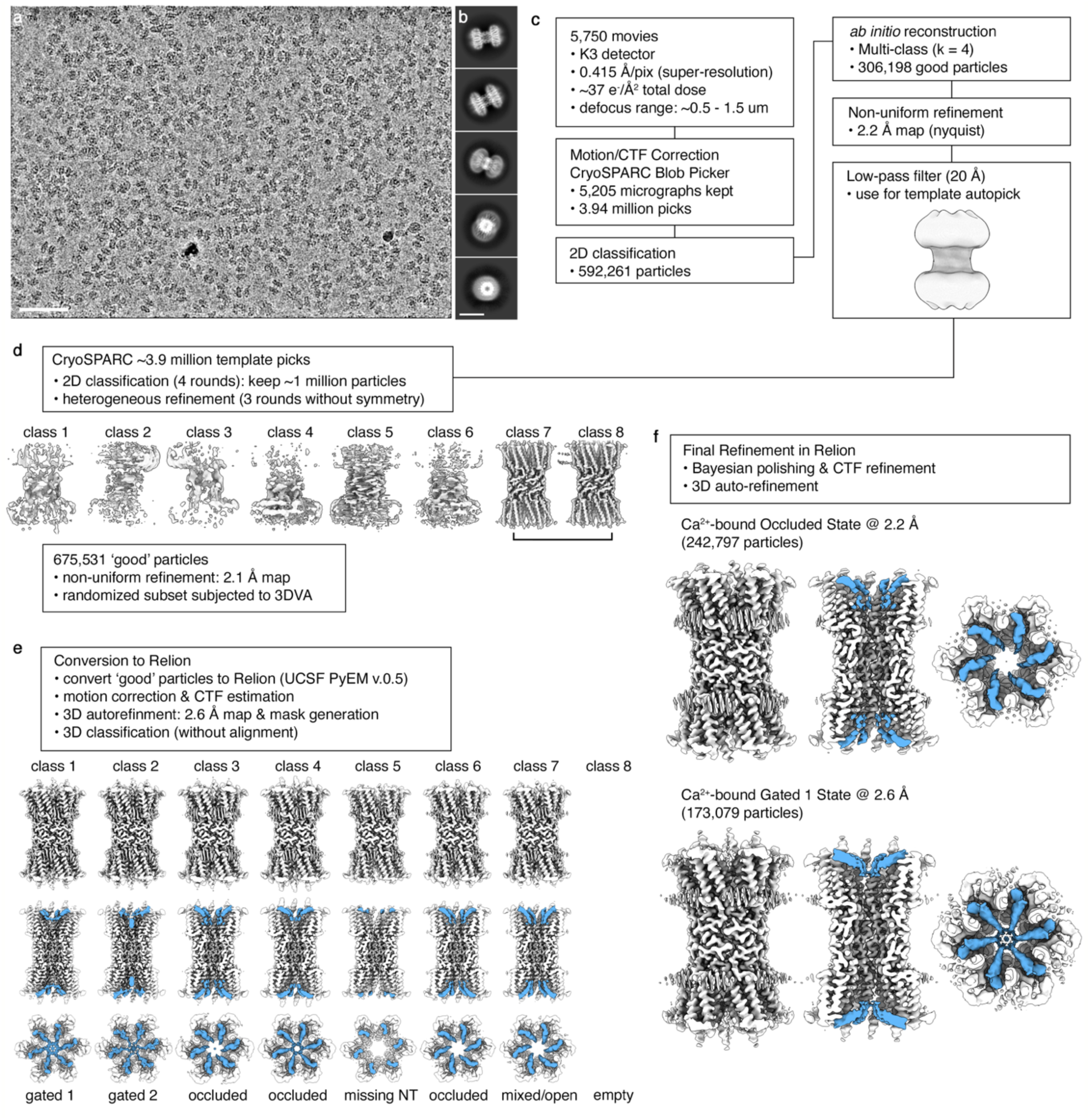
Cryo-EM image processing workflow. **a**, Representative cryo-EM micrograph from the 5,750 movie dataset recorded on a Gatan K3 detector. Scale bar = 50 nm. **b**, Representative 2D class averages. Scale bar = 10 nm. **c**, Image processing workflow to generate an initial reconstruction in CryoSPARC for 3D template picking. **d**, A combination of 2D classification and 3D heterogeneous refinement was used to clean up the template-picks to a dataset of 675,531 ‘good’ particles that refined to ∼2.1 Å resolution. A subset of these particles was subjected to symmetry expansion and 3D variability analysis (3DVA) in CryoSPARC. **e**, Particles were converted to RELION format for 3D classification, resolving the Ca^2+^-bound occluded, gated 1, and gated 2 states based on NT domain features (blue). Additional classes containing apparently mixed NT states, or missing NT density were excluded from further analysis. **f**, Per-particle polishing and 3D refinement produced ∼2.2 Å (Ca^2+^-bound occluded) and ∼2.6 Å (Ca^2+^-bound gated 1) maps suitable for atomic modeling. The Ca^2+^-bound gated 2 map lacked sufficient NT resolution for atomic-level interpretation.

**Extended Data Figure 3:**
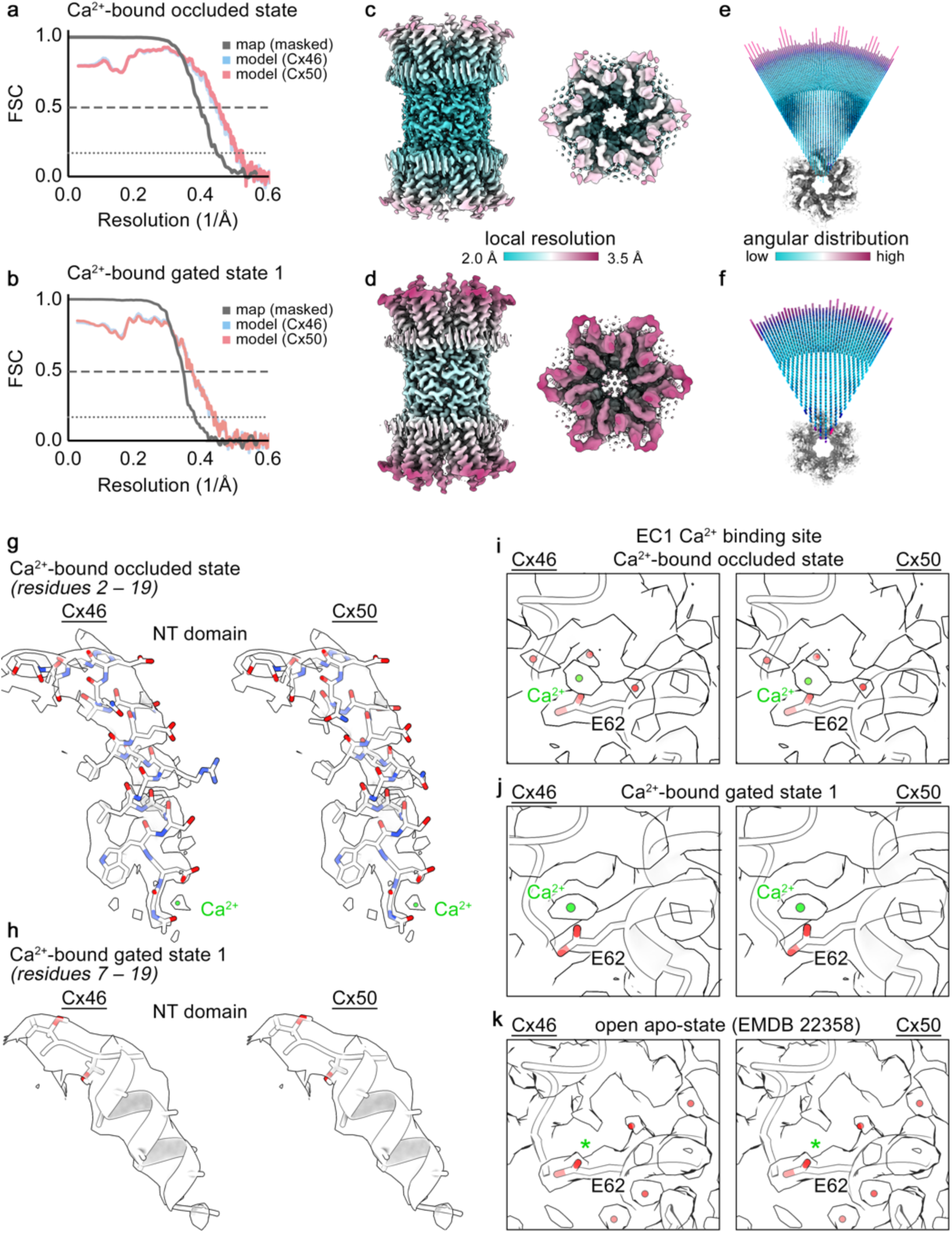
Global and local resolution assessments. **a-b**, Gold-standard Fourier shell correlation (FSC) for the Ca^2+^-bound occluded state and gated states. Half map correlation (black) and map-to-model (Cx50 – red; Cx46 – blue) shown, with cut-off values indicated (0.5 – dashed line; 0.143 – dotted line). **c-d**, Local resolution-filtered maps for the Ca^2+^-bound occluded and gated states, respectively, with resolution displayed by color gradient. **e-f**, Angular distribution plots for the occluded and gated states, with population occupancy indicated by color (magenta – high; cyan – low). **g-h**, Cx46 and Cx50 models fit to NT domains density for the occluded (residues 2–19) and gated (residues 7–19) states, respectively. Modeled Ca^2+^ ion (green) resolved at the NT site in the occluded state is labeled. Note, additional unassigned map densities near this site, limiting the assignment as putative. Sidechains for residues 7–17 of the gated state were truncated beyond Cβ due to limited local resolution. **i-j**, Zoomed views of the EC1 Ca^2+^ site in the occluded and gated states. Modeled Ca^2+^ ion (green), ordered water molecules (red), and E62 sidechain are shown. **k**, Corresponding region of the apo-state models fit to map density, with absence of density at the Ca^2+^ site indicated (asterisk).

**Extended Data Figure 4:**
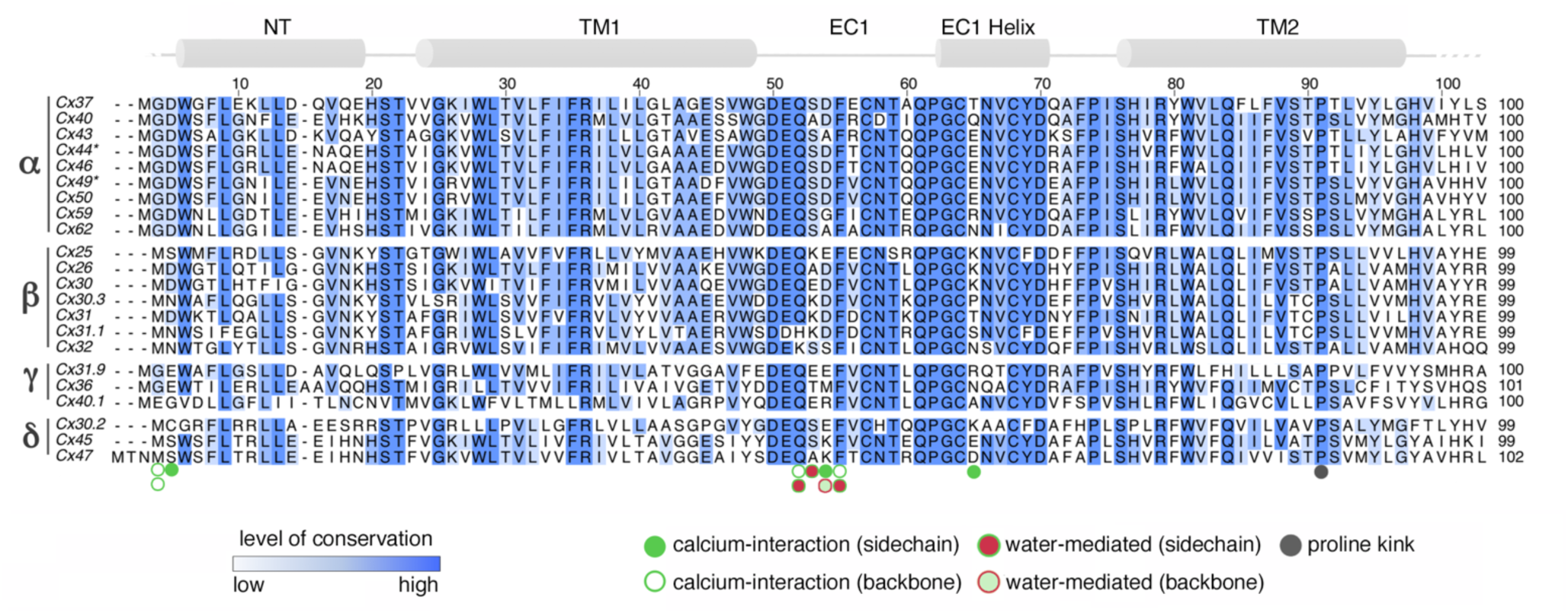
Sequence alignment of connexin pore-lining domains with Cx46/50 Ca^2+^ binding sites annotated. Multiple sequence alignment of 20 human connexin isoforms across the pore-lining regions (NT, TM1, EC1/EC1 helix, and TM2). Sheep homologs (Cx44 and Cx49, corresponding to Cx46 and Cx50, respectively) are included for comparison. Isoforms are grouped by connexin families (α, β, γ, δ); the orphan Cx23 is excluded. Regions of sequence conservation are indicated by the intensity of blue shading. Secondary structural elements, positions of Ca^2+^ binding sites, and proline kink are annotated, with Ca^2+^ interactions categorized by type (sidechain, backbone, or water-mediated) and color-coded as per the legend.

**Extended Data Figure 5:**
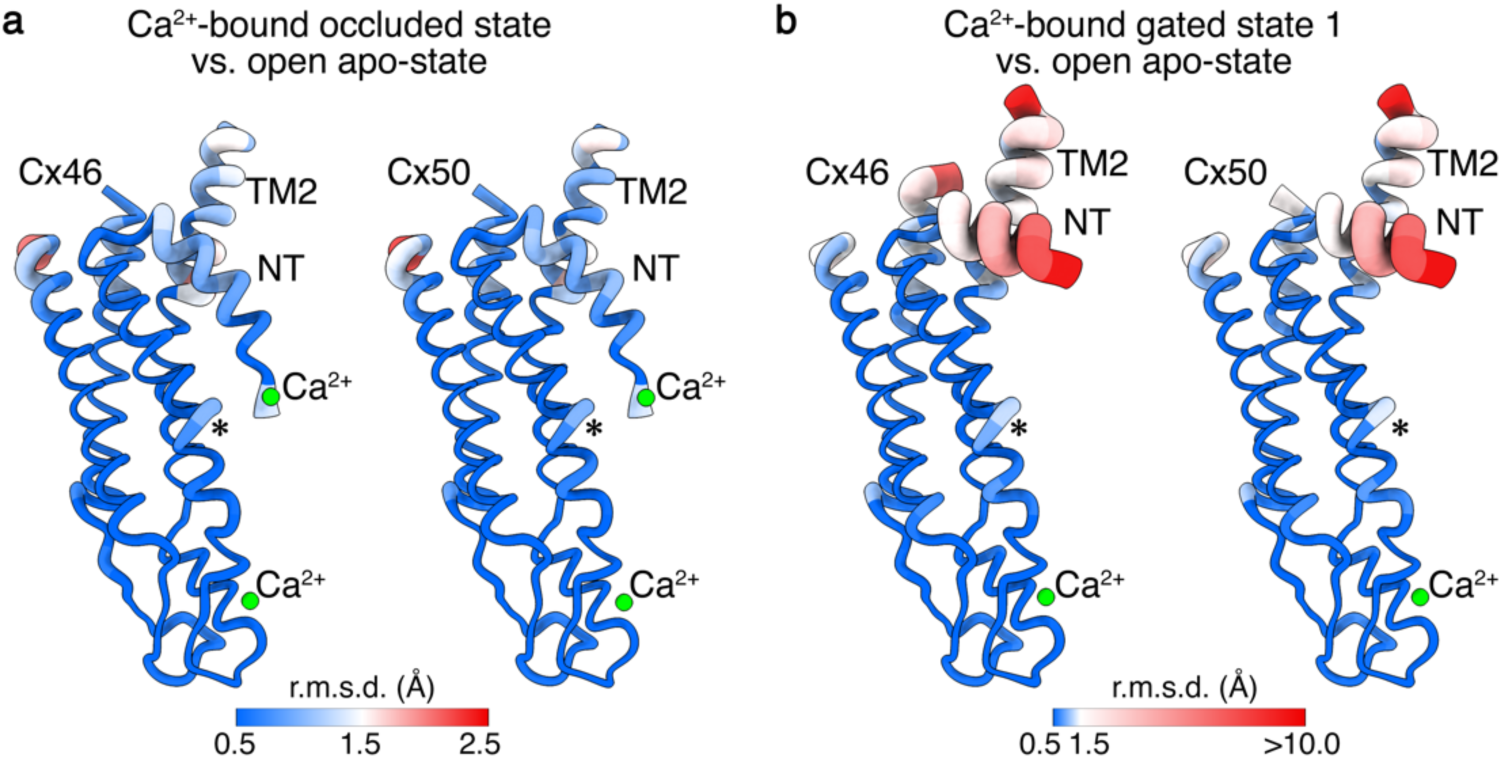
Structural comparisons of apo-state connexin-46/50 with Ca^2+^-bound states. Structural alignment and Cα root mean square deviation (r.m.s.d.) analysis of Cx46 and Cx50 in the open apo-state versus **a**, the Ca^2+^-bound occluded state and **b**, the Ca^2+^-bound gated 1 state. R.m.s.d. values are visualized by color gradient (blue to red, indicating low to high values) and by the thickness of the backbone in ‘worm’ representation produced in ChimeraX. NT, TM2, and Ca^2+^ binding sites are labeled. A key for r.m.s.d. values for each set of comparisons is displayed. Asterisk indicates region of minor variability around a π-helix that forms a kink in TM1 (residues ∼39-41).

**Extended Data Figure 6.**
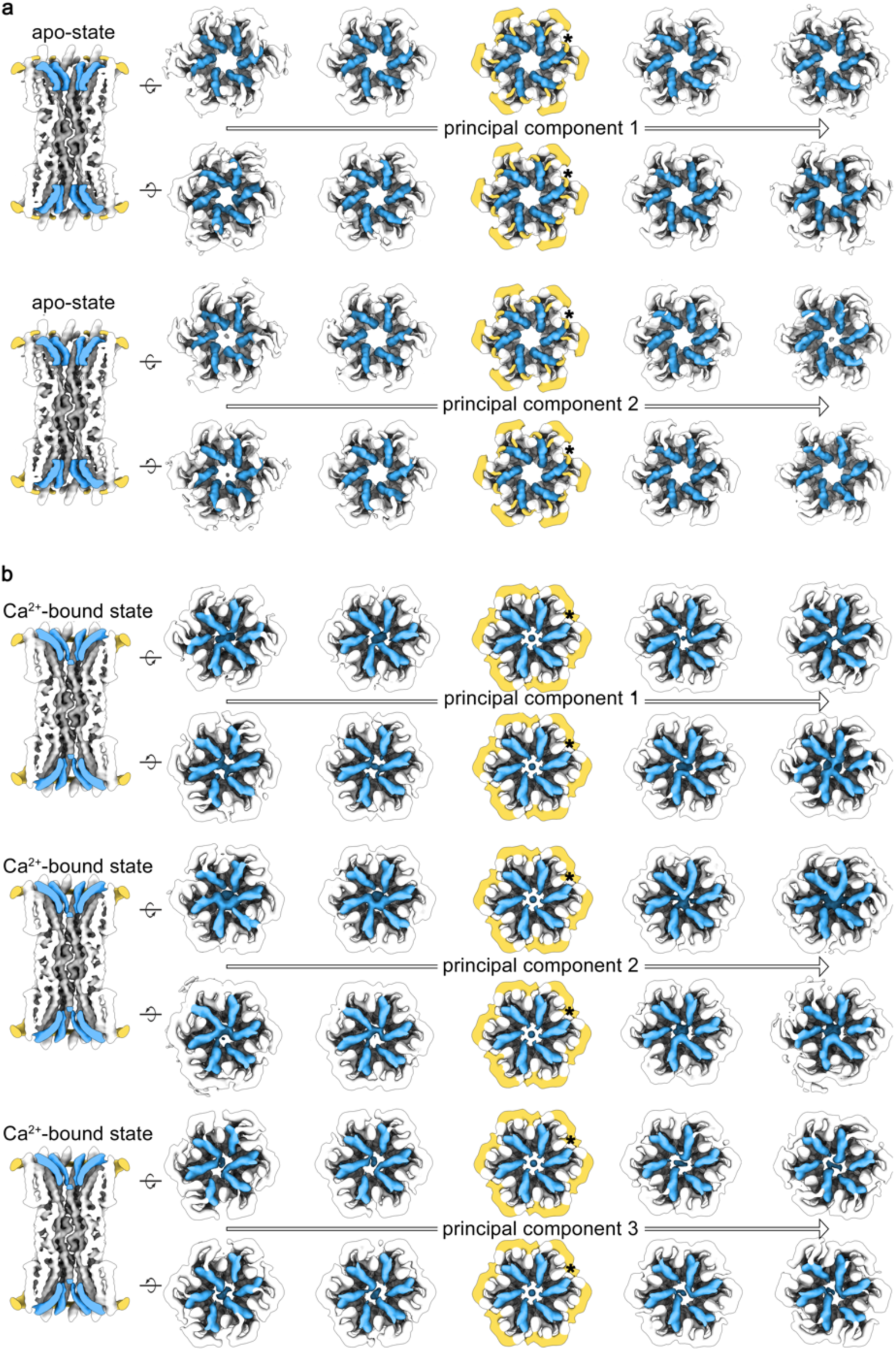
3D variability analysis of the apo-state and Ca^2+^-bound datasets. a,b, Representative frames from 3D variability analysis (3DVA) for the apo-state and Ca^2+^ bound datasets, respectively. A representative slice view from center of each principal component (PC) is shown (left), with corresponding ‘top’ and ‘bottom’ views (right). The NT domains (blue), ICL/CT domains (yellow) and TM2 (asterisk) are highlighted. For the apo-state dataset, the primary PC showed minimal variability, described primarily as a slight ‘wobble’ of the NT and TM2, with the pore remaining open. In the Ca^2+^ bound dataset, the first three PCs displayed significant variability in the NT, TM2, and ICL/CT domains. The NT domains displayed movement between occluded (or possibly open) and gated states, with coupling interactions between neighboring and/or opposing subunits within each hemichannel. The gated states are facilitated by pore-stretching, collapsing the pore distance to enable NT pairing and steric block of the pathway. TM2 orientation and ICL/CT reorganization correlated with NT movements.

## SUPPLEMENTAL MOVIE LEGENDS

- **Supplemental Movie 1. Morph between apo-state and Ca^2+^-bound occluded state.**
- **Supplemental Movie 2. Morph between apo-state and Ca^2+^-bound gated state.**
- **Supplemental Movie 3. 3D variability analysis (3DVA) of apo-state and Ca^2+^-bound datasets.**

## Notes

### Competing Interest Statement

The authors have declared no competing interest.

